# A role for class I PAKs in the regulation of the excitability of the actin cytoskeleton

**DOI:** 10.1101/2024.10.16.618394

**Authors:** Joe J. Tyler, Anthony Davidson, Megan E. Poxon, Montserrat Llanses Martinez, Pete Hume, Jason S. King, Vassilis Koronakis

**Author notes:** These authors contributed equally (Joe J. Tyler, Anthony Davidson).

## Abstract

P21 activated kinases (PAKs) are involved in a wide range of functions from the regulation of the cytoskeleton to the control of apoptosis and proliferation. Although many PAK substrates identified are implicated in the regulation of the actin cytoskeleton, a coherent picture of the total effect of PAK activation on the state of the actin cytoskeleton is unclear. We therefore set out to observe and quantify the effect of PAK inhibition on the actin cytoskeleton in greater detail. In Mouse Embryonic Fibroblasts, inhibition of PAK kinase activity, either by treatment with small molecule inhibitors or overexpression of mutant PAK constructs leads to the constitutive production of patches of the phosphoinositide PIP3 on the ventral surface of the cell. The formation of these patches remodels the actin cytoskeleton and polarises the cell. From the overexpression of truncated and mutant PAK constructs as well as an *in vitro* model of PAK recruitment to small GTPases we propose that this is due to a hyper recruitment of PAK and PAK binding partners in the absence of PAK kinase activity. This aberrant production of PIP3 suggests that, by limiting its own recruitment, the kinase activity of class I PAKs acts to downregulate PI3K activity, further highlighting class I PAKs as regulators of PI3K activity and therefore the excitability of the actin cytoskeleton.

## Introduction

The p21 Activated Kinases (PAKs) are a family of 6 serine/threonine kinases identified as acting downstream of the small GTPases Rac and Cdc42 (Edward Manser et al. 1994; 1995). The 6 family members are divided by sequence homology into two subclasses, class I (PAK 1,2,3) and class II (PAK 4,5,6) PAKs (Arias-romero and Chernoff 2008; Kumar et al. 2017). Whilst sharing many binding partners and substrates the PAK isoforms have different expression profiles and subcellular localisation patterns suggesting a separation of function. PAK substrates include proteins vital to the regulation of proliferation such as Raf1 (Chaudhary et al. 1999; Zang, Hayne, and Luo 2002), MEK1 (Parka, Eblenb, and Catling 2008), and β-catenin (G. Zhu et al. 2012) and as a consequence PAKs are often dysregulated during cancer progression and are considered important targets in the search for new therapeutics (King, Nicholas, and Wells 2014; Radu et al. 2014; Ong et al. 2011; Ye and Field 2012).

All PAK isoforms contain an N-terminal p21 Binding Domain (PBD) and a C-terminal kinase domain (Edward Manser et al. 1994) and adopt an autoinhibited state, mediated by an interaction between the kinase domain and the Auto Inhibitory Domain (AID) (Zhao et al. 1998). For class I PAKs this has been thought to be due to an intermolecular interaction that results in the formation of a homodimer (Lei et al. 2000; Baskaran et al. 2012). However recent evidence suggests that it may in fact be intramolecular, as is the case for class II PAKs (Sorrell and Kilian 2019). Regardless of the exact nature of the interaction, this autoinhibition is relieved by GTPase binding to the PBD domain.

Unsurprisingly for a kinase acting downstream of Rac/Cdc42, many PAK substrates are directly linked to the regulation of the cytoskeleton including LIM Kinase (LIMK) (Edwards et al. 1999), filamin (Vadlamudi et al. 2003), MLC Kinase (MLCK) (Sanders et al. 1999) and a component of the Arp2/3 complex, ArpC1B (Vadlamudi et al. 2004). Additionally, several kinase independent roles for class I PAKs in the regulation of the cytoskeleton have been identified (Higuchi et al. 2008; Davidson et al. 2021) and the transient overexpression of wt or mutant PAK constructs as well as the inhibition of PAK by small molecule inhibitors have all been shown to have dramatic consequences for the organisation of the actin cytoskeleton (Sells et al. 1997; Mary Ann Sells, Boyd, and Chernoff 1999; Dharmawardhane et al. 2000; M. Kim et al. 2009; E Manser et al. 1997; Itakura et al. 2013; Mierke et al. 2020). Despite the wide body of work linking PAKs to the regulation of the actin cytoskeleton, a cohesive picture of the exact role of PAKs in the regulation of the cytoskeleton remains elusive.

Since the average serine threonine kinase is predicted to phosphorylate 300 unique sites, this is perhaps unsurprising (Sugiyama, Imamura, and Ishihama 2019). However, the dysregulation of class I PAKs during cancer makes this an important issue, as changes to the actin cytoskeleton underpin many of the hallmarks of disease progression (Hideki Yamaguchia 2014; Olson and Sahai 2009). Having recently identified a kinase-independent role for class I PAKs during pathogen-mediated actin remodelling (Davidson et al. 2021) we wanted to take a more general look at the consequence of PAK kinase activity on the actin cytoskeleton. We find that, in Mouse Embryonic Fibroblasts (MEFs), the inhibition of PAK kinase activity drives a PI3K dependent rearrangement of the actin cytoskeleton. We show that this rearrangement depends upon the presence of PAK that lacks kinase activity but can bind to small GTPases and contains an intact AID domain. We hypothesise that this results from the maintenance of a kinase-independent scaffolding activity that is usually opposed by PAK autophosphorylation. This provides further evidence of complicated feedback mechanisms between PAK activation and PI3K activity and complicates the interpretation of experiments performed with either PAK kinase inhibitors or the overexpressed kinase dead PAK constructs.

## Results

### Pak kinase inhibition polarises MEFs

To investigate the consequence of PAK kinase inhibition we first observed Mouse Embryonic Fibroblasts (MEFs) before and after the addition of the ATP-competitive PAK kinase inhibitor G5555 by low magnification transmitted light microscopy **(Supp. movie1)**. Before the addition of the drug, cells appear relatively elongated and produce multiple small, short-lived protrusions. Upon the addition of G5555, MEFs rapidly polarise, producing broad, fan-like lamellipodia which completely alter the shape of the cell. To better characterise this transition MEFs were transiently transfected to express mApple-LifeAct to mark the actin cytoskeleton and imaged by widefield fluorescence microscopy before and following the addition of G5555. **(Figure1A)** is a montage taken from **(Supp. movie2)** which captures the transition. Before the addition of G5555 multiple small, short-lived, protrusions can be observed. Upon addition of the drug a new protrusion forms and steadily expands. This protrusion is relatively stable and dramatically alters the shape of the cell as it grows.

**Figure 1.**
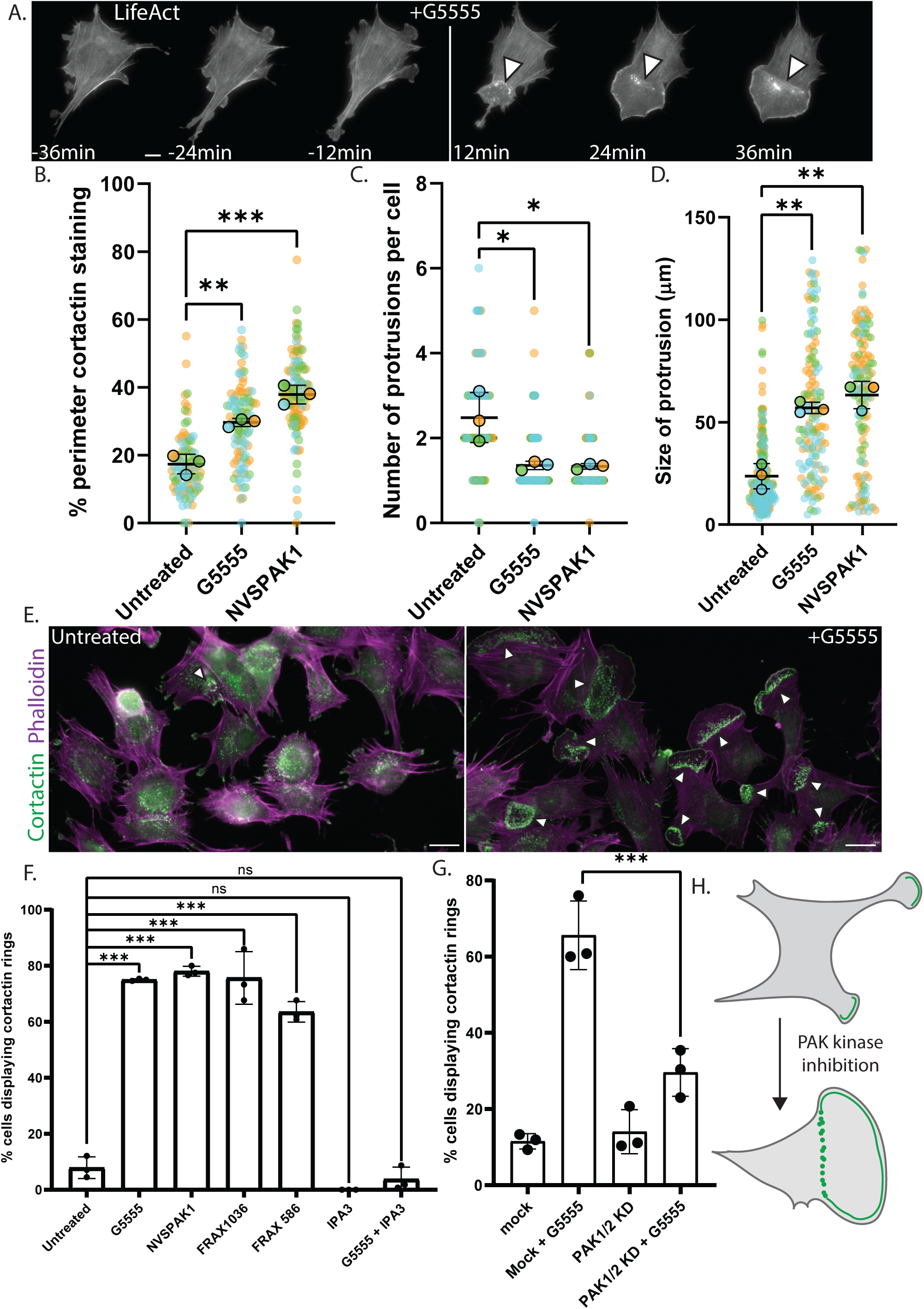
PAK kinase inhibition induces the formation of broad lamellipodia, polarising cells. A. Montage taken from movie of MEF transiently overexpressing mApple LifeAct, to mark the actin cytoskeleton, responding to the addition of G5555. Arrow indicates addition of G5555, arrow heads mark actin puncta at rear of newly formed protrusion. Scale bar indicate 10[gm B-D Quantification of changes in organization of the cytoskeleton and cell morphology upon PAK inhibition n=3 at least >100 cells analysed per condition total. B. Percentage of perimeter of cell marked by cortactin C. Number of protrusions per cell as defined by patches of cortactin staining at the periphery of the cell. D. Size of all individual protrusions analysed. E. Representative images highlighting upregulation of cortactin rings in the presence of G5555 scale bar 20µm. Arrow heads indicate cortactin/ actin ring (F) Percentage of cell producing cortactin rings in the presence of various PAK inhibitors. >100 cells analysed per condition per repeat. G. Effect of knockdown of PAK 1 and 2 expression on cortactin ring formation. H. Schematic representation of the consequence of PAK kinase inhibition in MEFs. All Error bars indicate SD. NS–no significant difference, *** -P ≤ 0.001 ** -P ≤ 0.01, * -P ≤ 0.05 (Ordinary one-way ANOVA followed by a post hoc Tukey’s multiple comparison test).

To quantify these observations MEFs were fixed and stained for cortactin, a marker of branched actin networks (Weed et al. 1999; Kage et al. 2022). Cortactin localisation to the periphery of the cell indicates an Arp2/3 dependent protrusion. Cells treated with G5555 showed an increase in the percentage of the perimeter that was marked by cortactin, demonstrating an increase in lamellipodia formation **(Figure1B)**. Whereas untreated cells produce multiple, small protrusions, Cells treated with G5555 produce on average, a single, large protrusion highlighting a polarisation in the presence of the drug **(Figure1C,D)**. A similar increase in lamellipodia production was observed in MDA-MB-231 and HAP cells, showing that this effect is not specific to MEFs **(Supp.1A,B,C,D)**. Furthermore, treatment of MEFs with the allosteric PAK kinase inhibitor NVSPAK1 (Karpov et al. 2015) resulted in a similar phenotype **(Figure1B,C,D)**. Together this suggests that the loss of PAK kinase activity promotes lamellipodia formation.

### PAK kinase inhibition drives formation of cortactin rings

Upon further analysis of **supp movie2** we noticed that the large protrusion that forms after addition of G5555 is marked by dynamic actin puncta which form a ring behind the leading edge **(arrowheads Figure1A)**. This further differentiates this structure from the smaller protrusions observed before the addition of G5555. To quantify the presence of these actin rings we re-examined images of cells fixed and stained for cortactin and found rings of cortactin positive actin puncta to be very prominent in cells treated with PAK kinase inhibitors **(Figure1E)**. Rings were clearly marked by cortactin, providing a convenient read out for the effect of the drug. Whilst observed in only 10% of untreated cells, 70-80% of cells treated with G5555 or NVSPAK1 display at least one ring of cortactin/ actin puncta **(Figure1F)**. A similar response was observed upon treatment with two additional ATP competitive Class I PAK kinase inhibitors FRAX 1036 and FRAX 597 (Semenova and Chernoff 2017). The inhibition of PAK kinase activity leads to the upregulation of a subset of cytoskeletal structures, typified by the presence of a ring of cortactin/actin puncta.

We took advantage of these cortactin/ actin puncta as a useful readout of the effect of PAK kinase inhibition to further characterise the effect of the loss of PAK kinase activity on the actin cytoskeleton. Like many kinases, several kinase-independent functions of class I PAKs have been previously identified (Davidson et al. 2021). Therefore, the kinase inhibitors tested so far may not block all functions of the molecule. IPA3 is a PAK inhibitor thought to block the GTPase binding activity of class I PAKs and therefore block recruitment (Deacon et al. 2008). To determine whether a kinase independent function of class I PAKs may contribute to the phenotype we have described, cells were treated with IPA3 and examined for the formation of cortactin/ actin rings. Interestingly IPA3 treatment alone did not drive the formation of cortactin/ actin whilst cotreatment of cells with G5555 and IPA3 blocked the effect of the kinase inhibitor **(Figure1E)**. This suggests that the recruitment of kinase-dead PAK is required for the observed changes to the cytoskeleton.

To examine this further, we partially depleted PAK1 and 2 by siRNA treatment and checked for the formation of cortactin/ actin rings **(Figure1G)**. Simultaneous knockdown of PAK1 and PAK2 had no effect on ring formation in untreated cells but did impair the ability of G5555 to induce cortactin/actin rings. This further suggests that the formation of cortactin/ actin rings in the presence of kinase inhibitors is due to the local recruitment of kinase dead PAK rather than a global reduction in total class 1 PAK activity.

### Effect of Kinase dead PAK requires GTPase binding and the AID domain

To determine how PAK kinase inhibition influences the cytoskeleton a series of truncated PAK constructs were expressed in MEFs, and these cells were assessed for the presence of cortactin rings. A schematic of PAK1 highlighting key interactions is shown in **(Figure2A)** and results of overexpression experiments are summarised in the table **(Figure2B).** In line with our siRNA results, simply increasing the global levels of full-length PAK1 in the cell, via the overexpression of full-length GFP-PAK1, had no effect on the formation of cortactin rings **(Figure2C)**. Overexpression of a truncated construct lacking the kinase domain (GFP-PAK1 Δkin) however led to a fourfold increase in ring formation. Furthermore, overexpression of a full-length, kinase dead, construct (GFP-PAK1 K298R) also promoted ring formation (Brown et al. 1996). Together, this provides further evidence that ring formation is dependent on the recruitment of PAK that lacks kinase activity.

**Figure 2:**
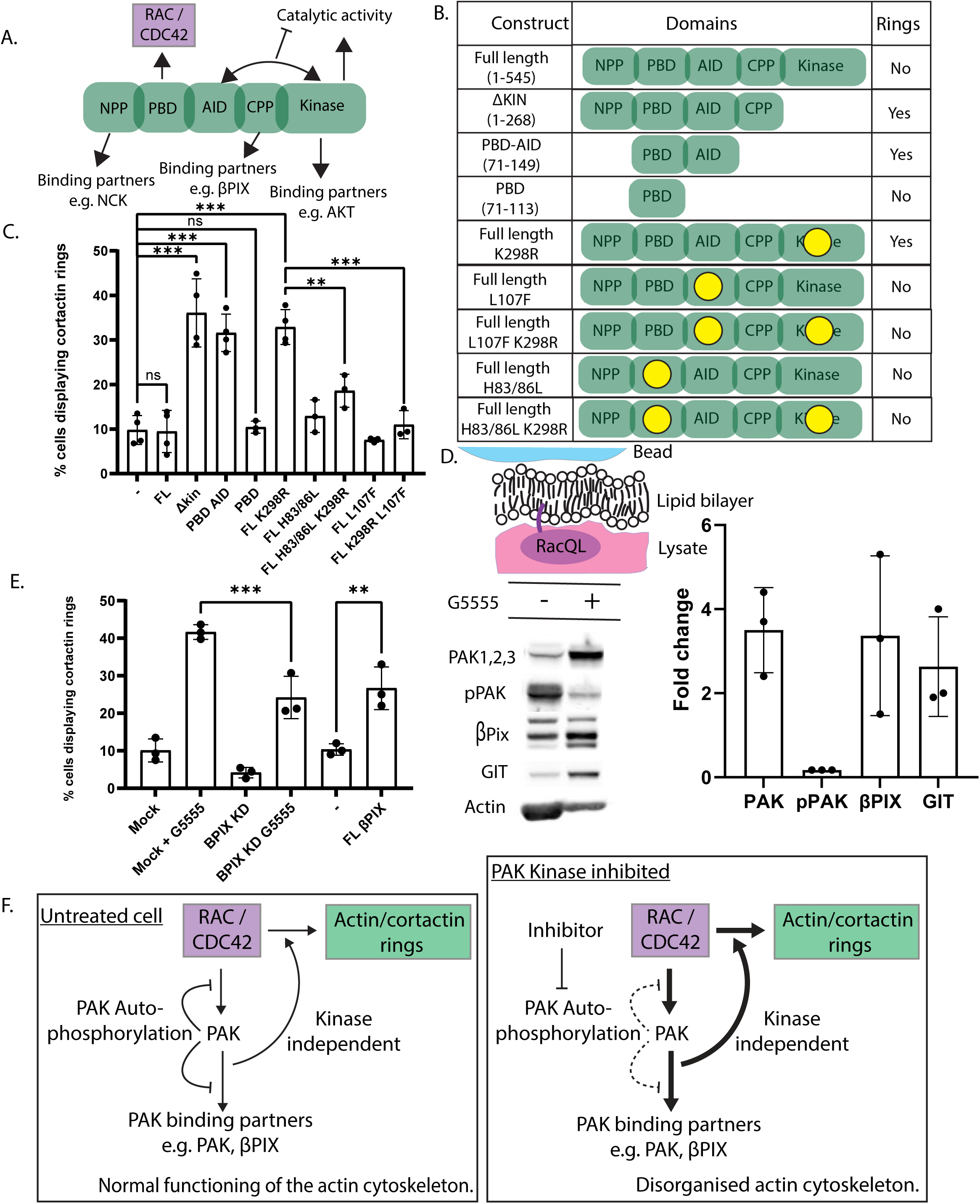
Overexpression of dominant negative PAK construct resembles treatment with PAK kinase inhibitors. A. Cartoon schematic indicating the domain organization of PAK and highlighting the localisation of a few key interactions. B. Table indicating identity of construct expressed and summarizing effect on cortactin ring formation C. Percentage of cells expressing various PAK constructs producing cortactin rings. n=<3 experiments each consisting of <100 cells. D. Schematic describing lipid coated bead experiment, western blot demonstrating levels of indicated protein associated to RacQL coated beads with and without addition of G5555 and quantification of fold change of indicated protein or phosphorylation event as detected by western blot. E. Percentage of cells producing cortactin rings in indicated conditions. n=<3 experiments each consisting of <100 cells. F. Schematic summary of proposed kinase dependent and independent effects on the formation of cortactin rings. All Error bars indicate SD. ns–no significant difference, *** -P ≤ 0.001 ** -P ≤ 0.01, * -P ≤ 0.05 (Ordinary one-way ANOVA followed by a post hoc Tukey’s multiple comparison test).

### GTPase-binding is required for cortactin/ actin ring formation

A further truncation consisting of the P21 Binding Domain (PBD) and Autoinhibitory Domain (AID) domain of PAK1 (GFP-PAK1 PBD-AID) promoted ring formation whilst a shorter construct consisting of just the PBD, regularly employed as a marker of active Rac, did not **(Figure2C)**. The PBD domain did however localise to the centre of actin rings in cells treated with G5555 **(Supp. 2a)**, suggesting that it may be sufficient to recruit PAK to these structures (and that Rac and/or CDC42 are active within the rings). To determine whether the GTPase binding activity of this domain contributes to the activity of kinase dead PAK, we overexpressed both a GTPase binding null full-length PAK1 (GFP-FL PAK1 H83/86L) and a GTPase binding null kinase dead FL PAK1 construct (GFP-FL PAK1 H83/86L K298R) and found that neither promoted ring formation **(Figure2C)**. The PBD is therefore sufficient to localise PAK1 to these structures, and both the GTPase binding activity of the PBD and the presence of the AID domain is required for the effect of kinase dead PAK on the cytoskeleton.

### Role of the AID domain

How the AID domain may contribute to the dysregulation of the actin cytoskeleton is unclear. The AID domain of PAK binds to the kinase domain, blocking the substrate-binding pocket and holding PAK in an autoinhibited state. To determine whether the interaction between the AID and kinase domain of PAK is required for the effect of kinase dead PAK we took advantage of the L107F mutation, which has previously been shown to disrupt this interaction (Brown et al. 1996; Frost et al. 1998). Overexpression of either GFP-full-length PAK1 L107F or GFP-full-length PAK1 L107F K298R did not promote the formation of cortactin/ actin rings **(Figure2C)** suggesting that the interaction between the AID and the kinase domain is required for the formation of these structures.

### Kinase dead PAK hyperaccumulates at active Rac

Since both the interaction with small GTPases and the ability to bind to more PAK is required for cortactin/ actin ring formation, we hypothesised that PAK kinase inhibition may lead to a change in the recruitment of PAK to active GTPases. Upon activation PAK undergoes a series of autophosphorylations which weaken both the interaction between the AID and the kinase domain (Chong et al. 2001) as well as the interaction between PAK and Rac (Edward Manser et al. 1994). In the absence of PAK kinase activity perhaps maintenance of tight GTPase binding, as well as the stabilisation of intermolecular PAK-PAK interactions, disrupts PAK localisation.

To test this, we used an *in vitro* model of PAK activation. Lipid coated beads coated with prenylated constitutively active Rac1 (Rac1 QL) were incubated with pig brain lysate, in the presence or absence of G5555 and protein recruitment to the beads was assessed by western blot **(Figure2D)**. In untreated lysates class I PAKs were pulled down by Rac1 QL and PAK phosphorylation at S144 was detected indicating PAK activation. Importantly this phosphorylation was lost in the presence of G5555 showing both that it is an active process occurring in the lysate and that the drug is working in this system. Upon inhibition of PAK kinase activity, 3 times more PAK was recruited to Rac1 QL beads. Loss of PAK kinase activity therefore leads to the aberrant accumulation of PAK, suggesting that one consequence of PAK kinase activity is to limit PAK recruitment to sites of small GTPase activation **(Figure2D)**.

### Role of βPIX during PAK inhibition

We next asked how accumulation of kinase dead PAK at small GTPases influences the organisation of the cytoskeleton. We hypothesised that enrichment of PAK may lead to the accumulation of PAK binding partners. βpix is a Rac and Cdc42 GEF that binds to PAK and is therefore a good candidate to mediate changes to the actin cytoskeleton (Edward Manser et al. 1998). Western blot analysis showed that both βpix and its constitutive binding partner GIT were enriched on RacQL beads in the presence of the PAK kinase inhibitor **(Figure2D)**. In agreement with a potential role for βpix during PAK kinase inhibition, partial depletion of βPIX by siRNA treatment led to a 50% reduction in the number of cells that generated cortactin/ actin positive rings upon treatment with G5555 **(Figure2E)**. To further confirm a role for βPIX in this process we transiently overexpressed βPIX -GFP and checked for cortactin/ actin ring formation. Upon overexpression of this construct there was a 2.5-fold increase in the number of cells making rings **(Figure2E).**

βPIX is required for the full response of the cell to PAK kinase inhibition and overexpression of full length βPIX alone is sufficient to mimic the phenotype of PAK inhibition. We therefore conclude that the dramatic rearrangements of the actin cytoskeleton upon PAK kinase inhibition depends upon the presence of a molecule of PAK that lacks catalytic activity but maintains the GTPase binding activity of the PBD domain and the kinase binding activity of the AID domain. This results in the hyperaccumulation of PAK and PAK binding partners and the dysregulation of the actin cytoskeleton **(Figure2F)**.

### Identity and dynamics of cortactin rings

#### Identity of cortactin puncta

To determine how the recruitment of a kinase dead molecule of PAK might influence the cytoskeleton we set out to better define the cortactin/ actin rings observed in **(Figure1E)**. First, we asked where PAK1 localised. Live cell Total Internal Reflection fluorescence microscopy (TIRFm) of cells expressing GFP-PAK1 and mApple LifeAct, treated with G5555, showed PAK1 to localise within the actin ring **(Figure3A)**. GFP-PAK1 also showed clear localisation to focal adhesions across the ventral surface of the cell.

**Figure 3.**
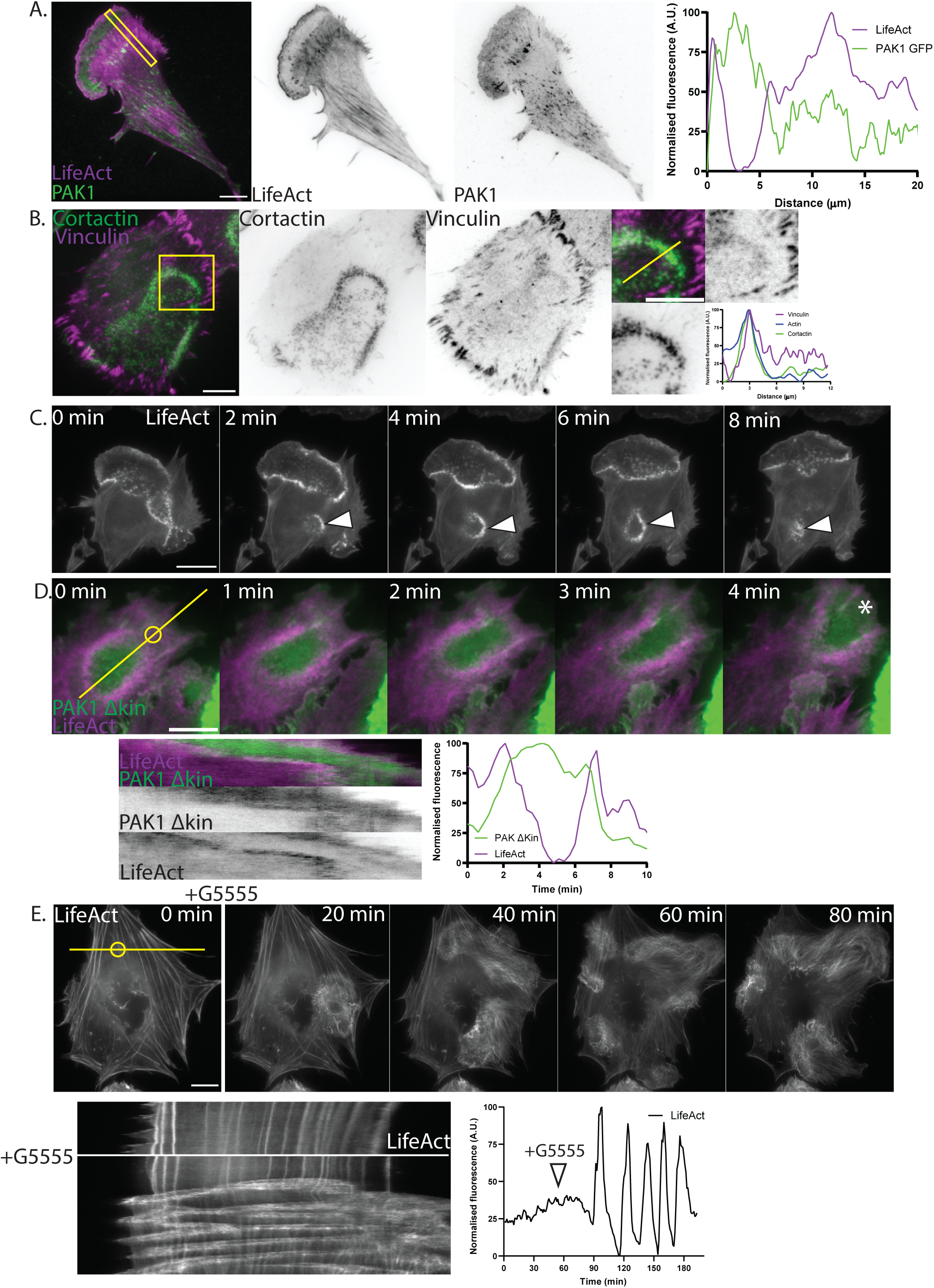
Identity and dynamics of cortactin rings. A. TIRFm image of a cell coexpressing GFP-PAK1 (Green) and maple LifeAct (magenta) demonstrating localisation of PAK1 to the centre of ring structure B. Representative image of cell stained with vinculin (green) and cortactin (magenta) antibodies showing localisation of vinculin to the ring of cortactin puncta. C. Montage taken from movie of cell transiently overexpressing mEmerald-LifeAct and treated with G5555 to highlight dynamics of internal rings D. Montage taken from supp movie of MEF expressing GFP PAK1 Δkin and mApple LifeAct. Accompanying kymograph taken from line indicated in the first panel. Graph indicates fluorescence intensity overtime in region indicated by circle in first panel. E. Montage taken from supp movie of B16F1 transiently overexpressing mEmerald-LifeAct responding to the acute addition of G5555. Accompanying kymograph generated from the same movie highlighting the disruption of the actin cytoskeleton upon addition of G5555 taken from line indicated in first panel. Graph indicates fluorescence over time in region of the cell marked by a circle in first panel. Scale bars 10µm unless otherwise stated.

Next, we further probed the molecular composition of the cortactin/ actin rings. A combination of live cell imaging of overexpressed fluorescent markers and immunostaining of endogenous proteins was used to dissect the molecular composition of the cortactin positive actin puncta. Unsurprisingly these were shown to contain several components of the Arp2/3 complex (**Supp. 3A)**. They were also marked by the nucleation promoting factor N-WASp **(Supp. 3B)** and two components of the wave regulatory complex (CYFIP and ABI) **(Supp. 3C,D)** as well as Myo1e (**Supp. 3E)**.

The size and molecular composition of these puncta are reminiscent of podosomes. These are actin dependent adhesive structures characterised by a dense actin core surrounded by a ring of adaptor proteins such as vinculin (Dries et al. 2019). Immunostaining of endogenous vinculin revealed a diffuse localisation around the actin puncta, suggesting that they may indeed be adhesive **(Figure3B)**. These puncta therefore appear similar in both superficial appearance and molecular composition to podosomes but lack the clear organisation of actin core and adhesive ring. We therefore tentatively term these structures ‘podosome-like’.

### Dynamics of cortactin rings

These ‘podosome like’ puncta are organised into higher order structures, referred to here as cortactin/ actin rings, that are themselves dynamic. To gain a better understanding of the identity of this superstructure we further examined time lapse movies of cells expressing mApple LifeAct following treatment with G5555 **(Supp. movie3** and **Figure3C)**. **Figure3C** highlights the dynamic nature of these structures. The ring of actin puncta can be seen to fluctuate in place; dramatically growing and shrinking. The actin in the centre of this ring appears different in organisation to the actin outside and where it contacts the edge of the cell it drives the formation of a protrusion.

Such dynamics are not typical of classic lamellipodia and are more reminiscent of those described for actin teeth in macrophages performing stalled phagocytosis or the wavefronts of actin observed during circular dorsal ruffle (CDR) formation (Barger et al. 2019; Ostrowski et al. 2019; Bernitt et al. 2015). Indeed smaller rings can be observed forming within and travelling across the body of the cell **(Arrows)**, similar to the actin waves observed in lab strains of *Dictyostelium* (Vicker 2002b; Bretschneider et al. 2009) as well as in certain mammalian cell types (Vicker 2002a; Zhan et al. 2020). These dynamics are further highlighted in **Supp. movie4** and **Figure3D**, a montage from TIRFm imaging of a cell expressing mApple-LifeAct and GFP-PAK1 ΔKin. Here a patch of PAK1 ΔKin is seen surrounded by a dynamic ring of actin. As the patch moves so does the ring, until it contacts the periphery of the cell and drives the formation of a protrusion (indicated by * in final panel). The correlation of the PAK1 ΔKin patch and actin ring is highlighted in the accompanying kymograph which was generated along the line indicated in the first panel of **Figure3D**. The accompanying graph shows fluorescence intensity over time within the region marked with a circle in the first panel of **Figure3D.** This further shows how the actin cytoskeleton remodels as the patch of kinase dead PAK travels across the ventral surface of the cell.

Finally, we wanted to determine whether similar dynamics could be observed in another cell line. B16F1 cells are routinely used in studies of the actin cytoskeleton for a number of reasons, not least because they form prominent lamellipodia (Kage et al. 2022; Buracco et al. 2022). We hypothesised that the apparent excitability of the cytoskeleton in this cell line might make for a prominent response to PAK kinase inhibition. B16F1 cells were therefore transiently transfected to express eGFP lifeAct and observed via widefield fluorescence microscopy before and then during the addition of G5555 **Figure3E, Supp. movie5**. Before addition of the drug the cytoskeleton of the cell appears relatively stable with prominent stress fibres and only small, short-lived protrusions. Following the addition of G5555 an actin wave can be seen to form that begins to traverse around the cell. This wave splits at multiple points until many waves can be seen travelling around the cell, driving protrusion where they contact the cell edge and reorganising the prominent stress fibres where they pass. The dramatic change in the state of the cytoskeleton is highlighted in the accompanying kymograph as well as a read out of actin intensity overtime at the region of the cell marked with a circle in the first panel.

### Lipid composition of rings

Actin waves and related structures are organised around regions of the plasma membrane with a specific identity, within which PI(3,4,5)P_3_ /PI(3,4)P_2_ are enriched and PI(4,5)P_2_ is depleted (Zhan et al. 2020; Masters, Sheetz, and Gauthier 2016; Brandon 2021; Ostrowski et al. 2019; Gerhardt et al. 2014). To examine whether a similar organisation of lipid species may organise the cortactin/ actin ring structures observed upon inhibition of PAK kinase activity, fluorescent markers of various phosphoinositides were expressed in MEFs expressing mApple LifeAct to mark the actin cytoskeleton. These cells were then treated with G5555 and the localisation of the different markers relative to the actin cytoskeleton compared using TIRFm.

After treatment with G5555 the PI(3,4,5)P3 marker (GFP BTK PH) was observed to localise within the ring of actin puncta **(Figure4A, Supp. movie6)**. BTK fluorescence dropped off sharply outside of the ring and there was no corresponding peak of PI(3,4,5)P3 observed relative to the actin puncta. Interestingly the PI(3,4)P2 marker (GFP TAPP1 PH) and a PI3P markers (EYFP p40phox px domain) also localized within the actin ring **(Supp. 4A,B)**. The PI(4,5)P_2_ marker (mCherry PLC PH) was clearly excluded from the center of the ring of actin puncta **(Figure 4B, Supp. movie7)**.

**Figure 4.**
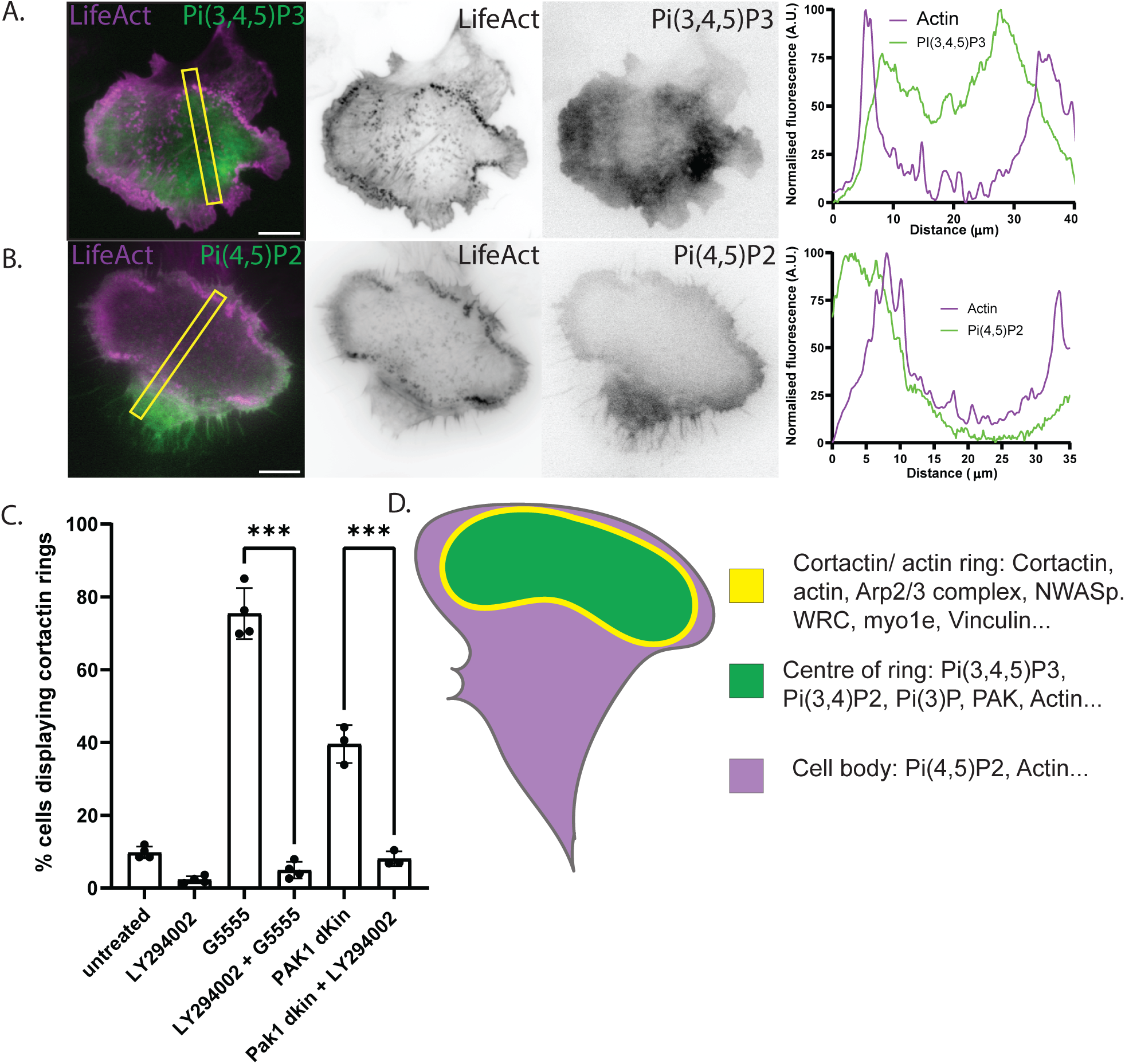
Formation of cortactin rings is PI3K dependent. A-B. Still from movie of MEF transiently overexpressing mCherry-LifeAct (magenta) and indicated phosphoinositide marker responding to the addition of G5555. Accompanying linescan demonstrates localization of lipid marker relative to actin cytoskeleton. C. Percentage of cell producing cortactin rings in the presence and absence of PI3K inhibitor LY294002. n=<3 experiments each consisting of <100 cells. D. Schematic summarising the localization of various proteins upon addition of G5555. All scale bars 10µm. All Error bars indicate SD. *** -P ≤ 0.001 (Ordinary one-way ANOVA followed by a post hoc Tukey’s multiple comparison test).

Cortactin/ actin rings therefore form around patches of membrane enriched in PIP3, PI(3,4)P2 and PI(3)P. Due to the prevalence of 3’ phosphorylated phosphoinositides within the centre of the rings we wanted to determine the role of PI3Ks on the formation of these structures. Cotreatment with the PI3k inhibitor LY294002 abolished cortactin ring formation in both cells treated with G5555 and cells expressing PAK1 Δkin **(Figure4C)**. The formation of these structures upon PAK kinase inhibition is therefore PI3K dependent. In agreement we detected an increase in the ratio of pAKT : AKT by western blot in cells treated with G5555, consistent with a global increase in PIP3 **(Supp. 4C)**. We propose then that the accumulation of PAK upon PAK kinase inhibition ultimately leads to a dysregulation of PI3K signalling and the constitutive production of PIP3 domains on the plasma membrane. These domains organise actin polymerisation resulting in the formation of rings of cortactin/ actin positive puncta which travel across the plasma membrane **(Figure4D)**.

### PI3K dependence of phenotypic changes upon PAK kinase inhibition

PIP3 is produced at the plasma membrane in response to acute external stimulation (Karunarathne et al. 2013). This is one mechanism by which the actin cytoskeleton may be organised in response to a change in the external environment of the cell. The aberrant production of PIP3 in the presence of PAK kinase inhibitors may therefore mimic the application of an external signal. This might explain the polarisation of the cytoskeleton in the absence of an external directional cue, described in **Figure1**. We therefore further analysed the localisation of cortactin in cells treated with LY294002 or cotreated with LY294002 and G5555. Whilst cells treated with LY294002 still produced lamellipodia, this blocked the increase in lamelipodia formation when co-treated with G5555 **(Supp. 5A)**. Furthermore, there was no change in the number of protrusions formed per cell demonstrating that MEFs treated with G5555 do not polarise in the absence of PI3K activity **(Figure 5A)**.

**Figure 5.**
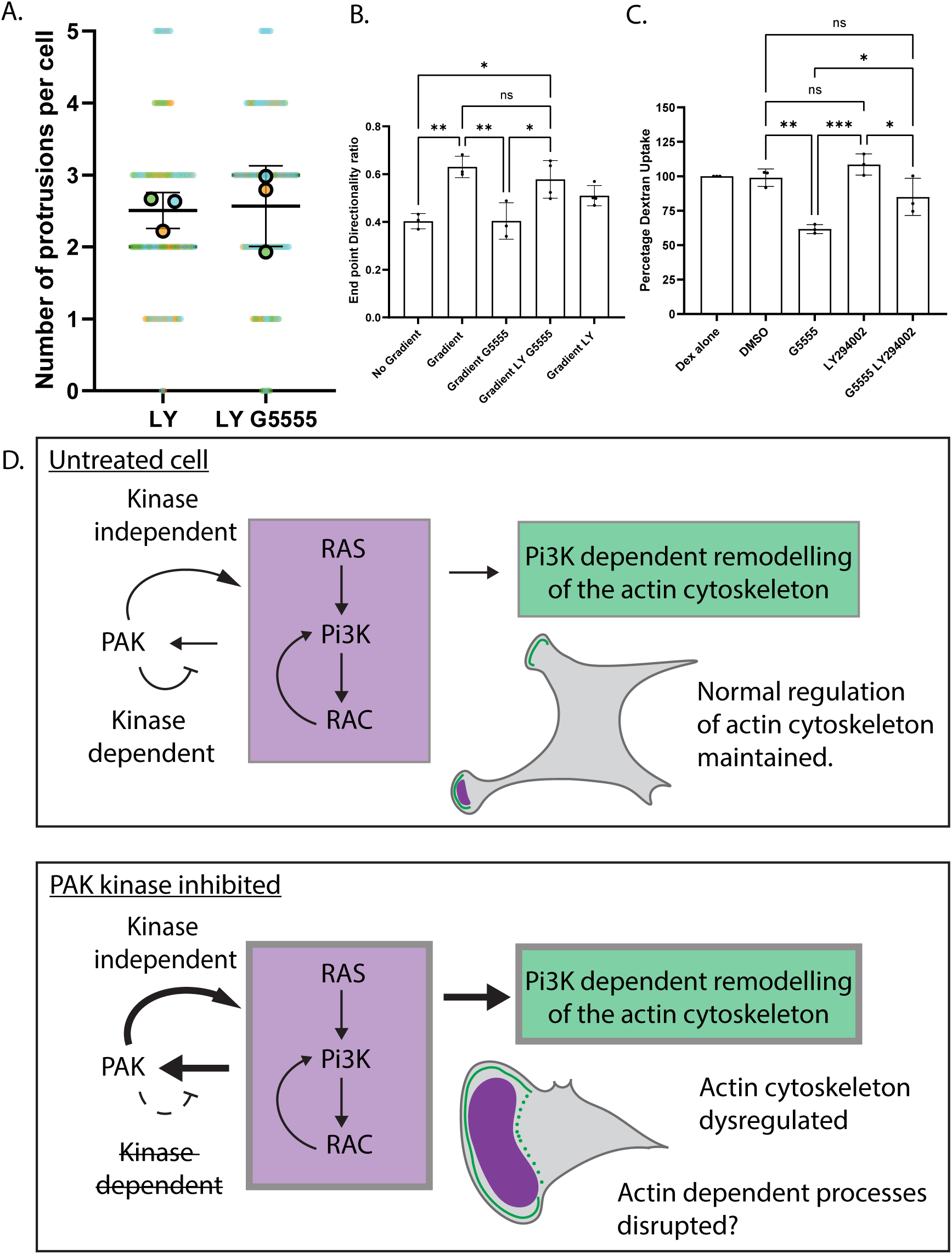
Pi3K dependence of phenotypic changes upon PAK kinase inhibition. A. Quantification of the number of protrusions per cell as defined by patches of cortactin staining at the periphery of the cell in cells treated with LY294002 or cotreated with LY294002 and G5555. B. End point directionality ratio of cells in indicated conditions tracked over a period of 18hrs C. Relative dextran uptake in MiaPacca cells in indicated conditions. D. Schematic summarizing proposed mechanism by which PAK kinase inhibition disrupts the regulation of the actin cytoskeleton.All Error bars indicate SD. NS–no significant difference, *** -P ≤ 0.001 ** -P ≤ 0.01, * -P ≤ 0.05 (Ordinary one-way ANOVA followed by a post hoc Tukey’s multiple comparison test).

### Migration

We hypothesised that this PI3K-dependent polarisation of the cytoskeleton may disrupt the ability of the cell to respond to external signals as the pathway may already be saturated. To test this, we analysed the effect of PAK kinase inhibition on cells migrating randomly and in the presence of a chemotactic gradient. Class I PAKs have previously been implicated in the regulation of cell migration with both overexpression of mutant constructs and inhibition with small molecule inhibitors have been shown to influence both the speed and directionality of migrating cells (Itakura et al. 2013; Mary Ann Sells, Boyd, and Chernoff 1999). The aberrant over-production of PIP3 has also been shown to disrupt efficient cell migration (Veltman et al. 2014).

Upon the addition of a chemotactic gradient untreated cells responded by orientating migration along the gradient. One read out of this change is an increase in the directional persistence, as cells turn less as their migration becomes directed **(Figure 5B)**. Upon the addition of G5555 however MEFs no longer increased directional persistence in the presence of a chemotactic gradient. However, cotreatment with LY294002 restored the ability of cells treated with G5555 to migrate in a more persistent manner within a chemotactic gradient. Plots displaying a random subset of cell trajectories are displayed in **(Supp. 5B)**. This suggests that the aberrant production of PIP3 in the absence of PAK kinase activity disrupts the ability of the cell to respond to external signals. This does not mean that the dysregulation of PI3K accounts for all effects of PAK in cell migration but demonstrates that directional migration is possible in the absence of PAK kinase activity, if PI3K activity is also inhibited.

### Macropinocytosis

We lastly wanted to determine whether the dramatic rearrangement of the cytoskeleton observed upon PAK kinase inhibition may disrupt other actin dependent processes. Macropinocytosis is the bulk uptake of extracellular fluid and requires an actin dependent reorganisation of the plasma membrane. Overexpression of various mutant PAK constructs, treatment with PAK kinase inhibitors and knockdown of PAK1 has previously been shown to influence levels of macropinocytosis (Lee et al. 2019; Dharmawardhane et al. 2000; Nazemi et al. 2024). We therefore wanted to determine whether the PI3K dependent rearrangement of the cytoskeleton may explain some of the effect of PAK kinase inhibition on this process.

Since macropinocytosis is PI3K dependent in many cell types we used MiaPaCa cells, which constitutively perform PI3K independent macropinocytosis (S. H. Kim et al. 2022). We first examined the response of MiaPaCa cells to PAK kinase inhibition and observed an increase in area upon treatment with G5555. Importantly this spreading was PI3K dependent **(Supp. 5C)**. We next examined the effect of PAK inhibition on macropinocytosis, as measured by the uptake of 70kDa dextran **fig5c**. As expected, untreated MiaPaCa cells performed macropinocytosis and this was unaffected by the addition of LY294002. Addition of G5555 however led to a nearly 50% reduction in uptake. Amazingly, this effect was partially reversed by cotreatment with LY294002. This further demonstrates that the aberrant formation of PIP3 in response to PAK kinase inhibition can disrupt the normal functions of the actin cytoskeleton **(Figure5D)**. Again this does not mean that dysregulated PI3K signalling accounts for all effects of PAK during macropinocytosis as several studies have shown an effect on macropinocytosis following a knockdown of PAK (Nazemi et al. 2024; Montserrat Llanses Martinez et al. 2024) but clearly demonstrates that caution is required when attributing phenotypes to the loss of PAK kinase activity when using PAK kinase inhibitors alone.

## Discussion

Previous studies have reported marked effects on the cytoskeleton upon both the overexpression of PAK and mutant PAK constructs as well as treatment with small molecule inhibitors. In this study we quantify and further characterise an increase in lamelipodia formation upon inhibition of PAK kinase activity in MEFs as well as in several other cell lines. We show that this correlates with an increase in the presence of a subset of PI3K-dependent actin structures. We conclude that PAK kinase inhibition results in a dysregulation of Pi3K activity and that this explains a number of the phenotypic changes observed upon PAK kinase inhibition. We hypothesise that this is due to the disruption of a class I PAK kinase dependent negative feedback loop which in untreated cells limits the recruitment of PAK and PAK binding partners to sites of GTPase activation. Together this provides further evidence of feedback from class I PAK kinase activation to PI3K activity and positions PAK and PAK binding partners as vital nodes in the regulation of the excitable pathways that control PI3K signalling.

Numerous studies have demonstrated effects on the actin cytoskeleton upon overexpression of various mutant PAK constructs as well as inhibition of PAK kinase activity by small molecule inhibitor. We find that the formation of PI3K dependent cortactin/ actin rings only occurs when PAK kinase activity is inhibited. Importantly we show that kinase inhibition by small molecule inhibitors recapitulates the effect of the overexpression of kinase dead PAK constructs. This suggests that the effect of kinase dead constructs represents the genuine inhibition of endogenous PAK kinase activity and demonstrates that similar phenotypes can be driven at endogenous levels of PAK expression.

The model proposed in **fig2F** suggests that: 1. the recruitment of PAK and PAK binding partners modulates the actin cytoskeleton in a kinase-independent manner and that 2. PAK kinase activity limits the total accumulation of PAK to sites of small GTPase activation. In the presence of PAK kinase activity, recruitment of PAK to small GTPases is therefore self-limiting and an optimal level of PAK recruitment, and therefore PAK scaffolding activity, is achieved. It is interesting to speculate that levels of PAK recruitment may be optimally tuned to elicit an appropriate cytoskeletal response following small GTPase activation. It will be important in the future to better characterise this scaffolding activity in the presence of PAK kinase activity. For example, in the absence of PAK as a scaffold, are cells in which levels of class I PAKs have been lowered by knockdown or knockout able to organise a coherent response to an external signal?

The mechanism linking hyper recruitment of PAK and PAK binding partners to PI3K activation also requires further investigation. We show that βPIX is required for the full response of the cell to PAK kinase inhibition, but how βPIX mediates this response is uncertain. Furthermore, class I PAKs and the PIX-GIT complex interact with multiple components of the excitable pathway that regulates PI(3,4,5)P3 production including binding directly to PI3K (Papakonstanti and Stournaras 2002), Rac (Edward Manser et al. 1994), ERK (Yin et al. 2005) and AKT (Higuchi et al. 2008) as well as phosphorylating the Ras effector Raf1 (Chaudhary et al. 1999) and the tumour suppressor Merlin (Xiao et al. 2002). The consequences of these varied activities and how they are integrated into a coherent response upon PAK recruitment and activation is far from clear. Due to the complex nature of interactions accounted for by the PAK-PIX-GIT-complex, the accumulation of these proteins is likely to be complicated and perhaps non-linear (J. Zhu et al. 2020; Premont et al. 2004).

Regardless of the exact mechanism, in MEFs the loss of PAK kinase activity results in the production of ventral PIP3 domains implying a dysregulation of the excitable networks which regulate PI3K activity. Acute perturbations to this pathway have previously been shown to drive dramatic switches in cell shape in both *Dictyostelium* and mammalian cell lines in agreement with the polarisation observed in MEFs (Miao et al. 2017; 2019; Devreotes et al. 2017; Zhan et al. 2020; Trogoff et al. 2023; Kuhn et al. 2024). This positions PAK as a potential node at which the excitable pathways controlling PI3K activity may be regulated. Again, it will be important now to test whether the opposite is true: do increased levels of PAK kinase activity negatively regulate the production of PIP3? Partial depletion of PAK 1 and 2 by siRNA did not in itself promote the formation of cortactin/ actin rings but it would be interesting to test this further in a system with higher basal PI3K activity.

Pip3 domains organise the cytoskeleton across large regions of both time and space and facilitate macropinocytosis and phagocytosis as well as a subset of protrusions in a range of cell lines (Veltman et al. 2016; Lutton et al. 2023; Ostrowski et al. 2019). The presence of these structures has been correlated with oncogenic transformation (Zhan et al. 2020) and they have recently been implicated in the regulation of glycolysis (Zhan et al. 2024). They have been studied extensively in *Dictyostelium* as lab strains constitutively produce ventral PIP3 domains which are amenable to imaging via TIRFm (Gerhardt et al. 2014). Analogous ventral structures can be induced in a range of mammalian cell types, however, comparison between Mammalian and *Dictyostelium* models has been somewhat limited by the relative difficulty of culturing and transfecting the specialised cell lines required as well as due to the transient nature of the response of the mammalian system to stimulation. The dramatic and long-lasting induction of ventral PIP3 domains in MEFs and B16F1s therefore provides a convenient model to better characterise the organisation and dynamics of these structures in mammalian cell lines.

Lastly, the rapid formation of large pip3 domains following the inhibition of PAK kinase activity complicates the interpretation of results obtained in the presence of PAK kinase inhibitors. The formation of PIP3 at the plasma membrane necessarily influences many signalling pathways (Swanson, Yoshida, and Swanson 2019) and PIP3 domains organise many key components of the cytoskeleton, potentially limiting the availability of key factors. Furthermore, they dramatically remodel the cytoskeleton, specifically the ventral cortex. The formation of these structures, which at times encompass the majority of the ventral surface of the cell, necessarily influences the mode of adhesion as well as the contractility of the cell. This in turn may further influence the state of key signalling pathways as well as the physical properties of the cell itself. Together this makes it hard to uncouple direct effects of the loss of PAK kinase activity from indirect effects stemming from the modulation of PI3K signalling and provides further justification for the recent development of a PAK1 selective degrader (Chow et al. 2023).

## Methods

### Cell culture

MEFs, B16F1s, MiaPaCas and MDA-MB-231 were maintained in Dulbecco’s Modified Eagle Medium (DMEM) supplemented with 10% FBS and 100U/ml penicillin-streptomycin. HAPs were maintained in Iscove’s Modified Dulbecco’s Medium (IMDM) supplemented with 10% FBS and 100U/ml penicillin-streptomycin. Cells were maintained at 37°C and 5% CO_2_. All experiments were performed on glass coated with fibronectin (Gibco) following manufacturers instruction.

Transfections were performed via electroporation using the Neon system (Invitrogen) According to manufacturer’s instruction using approximately 3μg of plasmid per transfection. Transfection was performed 18 hours prior to the start of experiments. Small interfering RNA (siRNA) Knockdown of genes of interest were performed using transfection via Oligofectamine (Life-Technologies) according to the manufacturer’s instruction. Transfection was performed 72hr prior to the start of experiments. Knockdowns were performed using RNAs previously described in (Davidson et al. 2021), purchased from Qiagen.

Drug treatments were added 90 minutes prior to fixation and preparation for imaging. Unless otherwise stated, live cell imaging of cells responding to treatment was performed at least 1 hour after the addition of a drug. Imaging of the acute addition of a drug was performed by careful addition to the microscopy dish. All small molecule inhibitors were used at a final concentration of 10µM except for IPA3, which was used at 50µM. All small molecule inhibitors were acquired from Tocris.

For Immunofluorescence experiments cells were fixed in 4% PFA before permeabilisation in PBS with 0.1% tween 20. Blocking was performed in PBS with 5% BSA for one hour and primary antibodies applied overnight at 4°C, in PBS with 5% BSA. Proteins were visualised by addition of fluorescent secondary antibodies (Invitrogen) and the actin cytoskeleton visualised by addition of Texas-red Phalloidin (Life Tech).

### Plasmids

Most PAK and βPIX constructs were used previously in (Davidson et al. 2021). PAK PBD (aa71-113) was subcloned from FL Em-PAK1 as a template. FL GFP-PAK1 (H83/86L K298R) was made via site directed mutagenesis using FL Em-PAK1 (H83/86L) as a template. All other plasmids were acquired from addgene: mApple LifeAct (addgene plasmid #54747), mEGFP LifeAct (addgene plasmid #54610), mcherry NWASp (addgene plasmid #55164), Myo1e Mcherry (addgene plasmid #27698), Emerald-Vinculin (Addgene plasmid #54303), mCherry Arp2 (Addgene plasmid #54980), GFP BTK PH (Addgene plasmid #51463), PH domain of PLCdelta1 (Addgene plasmid #36075), GFP-PH-Tapp1 (#161985), p40PX-EYFP (addgene plasmid #19011), Cyfip GFP (Addgene Plasmid #109139).

### Antibodies

Rb anti human Cortactin GTX GTX113681. Ms anti PAK1/2/3 Santa Cruz Biotechnology sc-166887, rb anti human GIT1 Cell Signalling #2919, Rb anti P-PAK1,2,3^Ser144/141^ Abcam [3H12], Ms anti β-PIX Santa Cruz Biotechnology sc-393184, rb anti Vinculin Cell Signalling #4650, Alexa Fluor® 488 anti-human CD29 Antibody BioLegend #303016.

### Microscopy

Live cell microscopy was performed at 37°C in CO_2_ independent media without phenol red (Leibovitz L15 medium without phenol red ThermoFisher). Fluorescence microscopy was performed on an Olympus IX83 inverted microscope equipped with an Olympus IX3-SSU automated xy stage and IX3 Z-Drift Compensator. Images were taken using CellSens software and recorded on a Hamamatsu imageEM x2 EM-CCD camera. TIRFm was performed with a Uapon 100x/ 1.49NA objective. Widefield fluorescence microscopy of live and fixed samples was performed on the same system with either 100x or 60x objective. Images of fixed MiaPaCa cells were acquired on Nikon W1 spinning disk with a 100x objective and images of mda-231 cells acquired with a Zeiss LSM980 Airyscan 2 with a 63x oil objective both within the Wolfson Light Microscopy facility at the university of Sheffield.

### Image analysis

All image analysis was performed in Fiji (Schindelin et al. 2019). To calculate the percentage of the perimeter of the cell marked by cortactin staining, first the total perimeter of the cell was acquired by manual segmentation. Next the length of each region of peripheral cortactin staining was measured. The cumulative total used to calculate the total percentage of cortactin staining. The number of protrusions per cell was given by the number of unique regions of peripheral cortactin staining per cell. Protrusion size was given by the length of all individual regions. The percentage of cells displaying cortactin/ actin rings was acquired by counting. A cortactin/ actin ring was defined as either a complete circle of dense cortactin/ actin staining within the body of the cell or an incomplete circle against the periphery of the cell, behind a protrusion (marked by a narrower band of cortactin/actin staining). Examples of both are visible in fig 1E.

### *In vitro* pull-downs with lipid coated beads

Pull-downs with lipid coated beads were performed as described previously (Hume, Humphreys, and Koronakis 2014). Briefly, silica microspheres were coated with phosphatidylcholine (PC) and phosphatidylserine (PS) at a molar ratio of 80:20 (Avanti Polar Lipids). These microspheres were incubated with prenylated RacQL for 2 hours before washing. This process is described in detail in (Hume, Humphreys, and Koronakis 2014). Beads were then incubated in cell free porcine brain extract for 30 minutes before extensive washing. As indicated, G5555 (10μM) or equivalent volume of DMSO alone was added to extract immediately prior to addition to beads. Recruited proteins were detected by western blot analysis. Blots were imaged on a LI-COR Odyssey Fc imaging system and band intensities quantified using Image Studio^TM^ software (LI-COR).

### Macropinocytosis

24 hours prior to analysis, cells were seeded at a density of 1.5 x 10^5^ cells in 200 μl/well of complete medium into flat-bottom, 96-well plates (ThermoFisher, 168055) each condition was seeded in duplicate, and incubated at 37°C. On the day of analysis, cells were pre-incubated with the respective inhibitors for 30 minutes, after which the drugs were co-incubated with 0.25 mg/mL lysine fixable Tetramethylrhodamine (TMR) conjugated 70 kDa dextran (ThermoFisher, D1818) for 90 minutes at 37 °C. After incubation, cells were washed 3 x with cold PBS, 100 μl of Trypsin was added to detach cells, followed by 100 μl of complete medium to neutralise the Trypsin. Cells were in fixed in 50 uL/well 4% paraformaldehyde (PFA) to a final concentration of 1%. Cells were analysed by flow cytometry using the YL1 laser on an Attune NxT Flow Cytometer (ThermoFisher) using the high-throughput sampling attachment, which pipetted them up and down twice, before analysing 150 μl per sample at 100 μl/second. At least 10,000 cells were counted per sample. Data was then inputted to GraphPad prism and the mean of three biological replicates plotted.

### Chemotaxis

Chemotaxis assays were performed in Ibidi µ-Slide Chemotaxis chambers according to manufacturer’s instructions. Chemotaxis was observed by transmitted light microscopy at 10x magnification on a Nikon ix83 inverted microscope at 37°C and cells were imaged every 5 minutes for 16.5 hrs. Cells were manually tracked using the manual tracking plugin in fiji. End point directionality, speed, and direction autocorrelation was calculated in Microsoft excel using DiPer as described in (Gorelik and Gautreau 2014). Plots demonstrating individual tracts were also generated in DiPer.

## Acknowledgments

This work was supported by grants to VK from the Welcome Trust (Grant number: 101828/Z/13/Z) and MRC (Grant number: MR/V000616/1) as well as the BBSRC (BB/W006049/1). We are grateful to the Wolfson Light Microscopy Facility for use of the Nikon W1 spinning disk confocal (BB/V019368/1) and Zeiss LSM980 Airyscan 2 Confocal (MR/X012077/1).

