## Supplemental Figures for "A role for class I PAKs in the regulation of the excitability of the actin cytoskeleton"

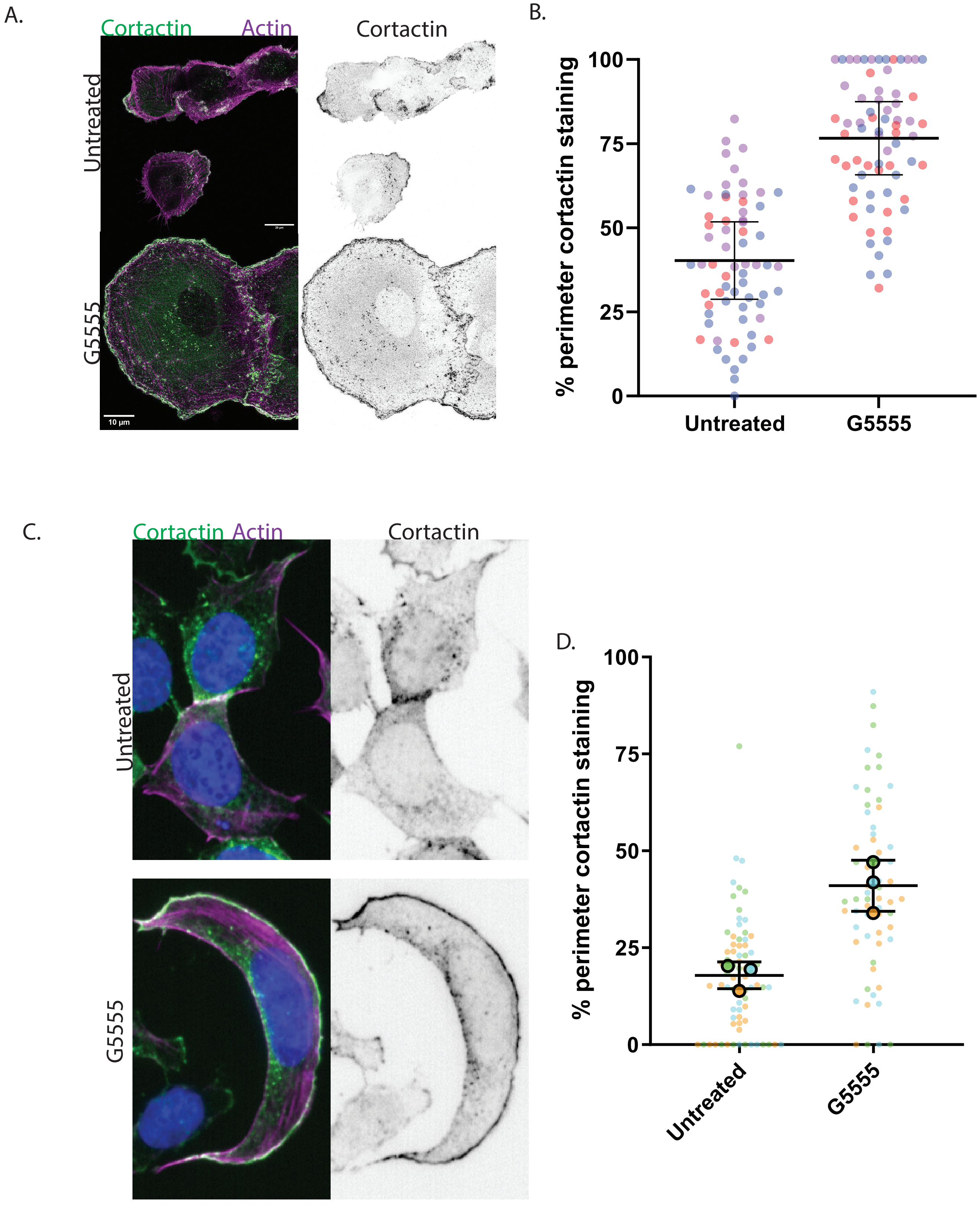


Supplemental Figure 1: PAK inhibition in MDA-MB-231s and HAPs. A. Representative images highlighting upregulation of cortactin rings in the presence of G5555 in MDA-MB-231 cells scale bar 10µm. B. Quantification of the percentage of perimeter of cell marked by cortactin with or without PAK kinase inhibition. C. Representative images highlighting upregulation of cortactin rings in the presence of G5555 in HAP cells. D. Quantification of the percentage of perimeter of cell marked by cortactin in HAP cells with or without PAK kinase inhibition.

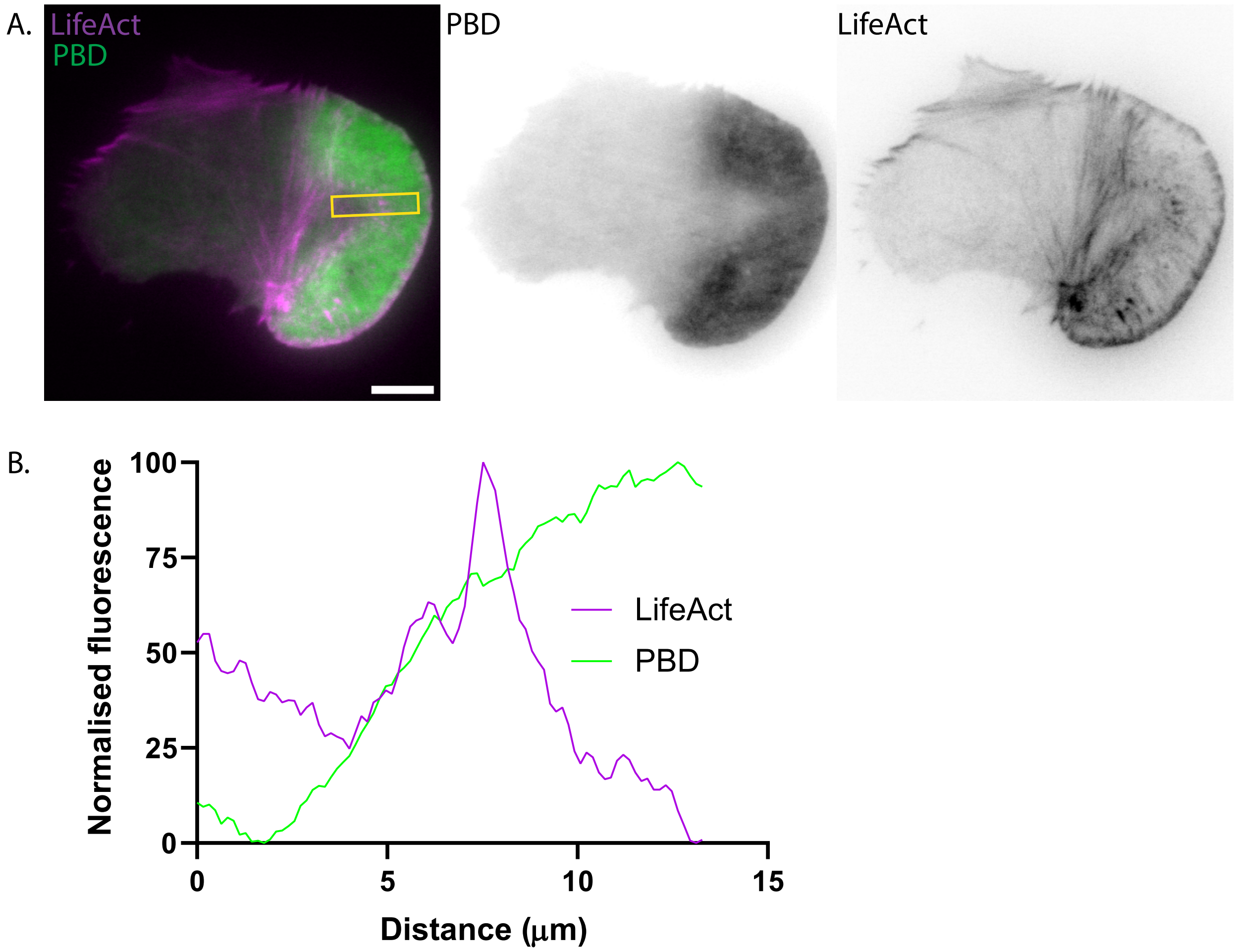


Supplemental Figure 2: A. PBD domain sufficient to localise PAK to actin ring structures. A. Representative TIRFm image of MEF expressing LifeAct (Magenta) and GFP-PAK PBD following treatment with G5555. B. Line scan taken along position indicated in Supplemental Figure 2A highlighting localisation of GFP-PAK PBD relative to peak of LifeAct signal that represents the edge of the actin ring structure. scale bar 10µm

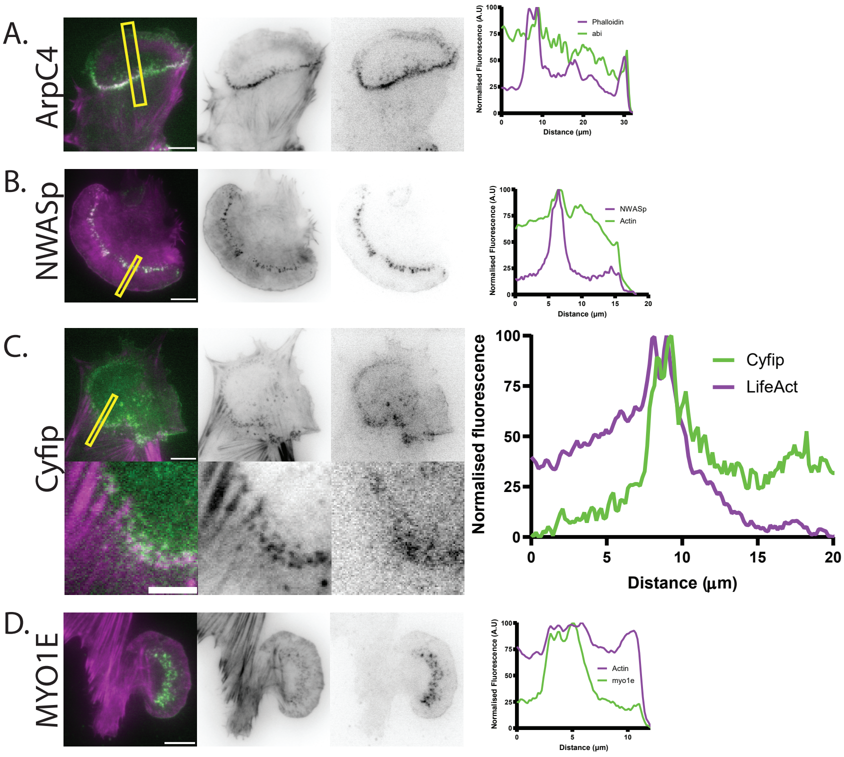


Supplemental Figure 3: Localisation of actin regulators relative to actin ring. A-D. Representative TIRFm image of cell expressing LifeAct (shown in magenta) and indicated fluorescently tagged actin regulator (shown in green), treated with G5555. Accompanying linescan highlights localisation of regulator to actin ring. A. MEF expressing mcherry-ArpC4 and mEGFP-LifeAct. B. MEF expressing mCherry-N-WASp and mEGFP-LifeAct. C. MEF expressing Cyfip-GFP and mCherry-LifeAct. Scale bar in magnified panel 5µm. D. MEF expressing Myo1e-mCherry and mEGFP LifeAct. All scale bars 10µm

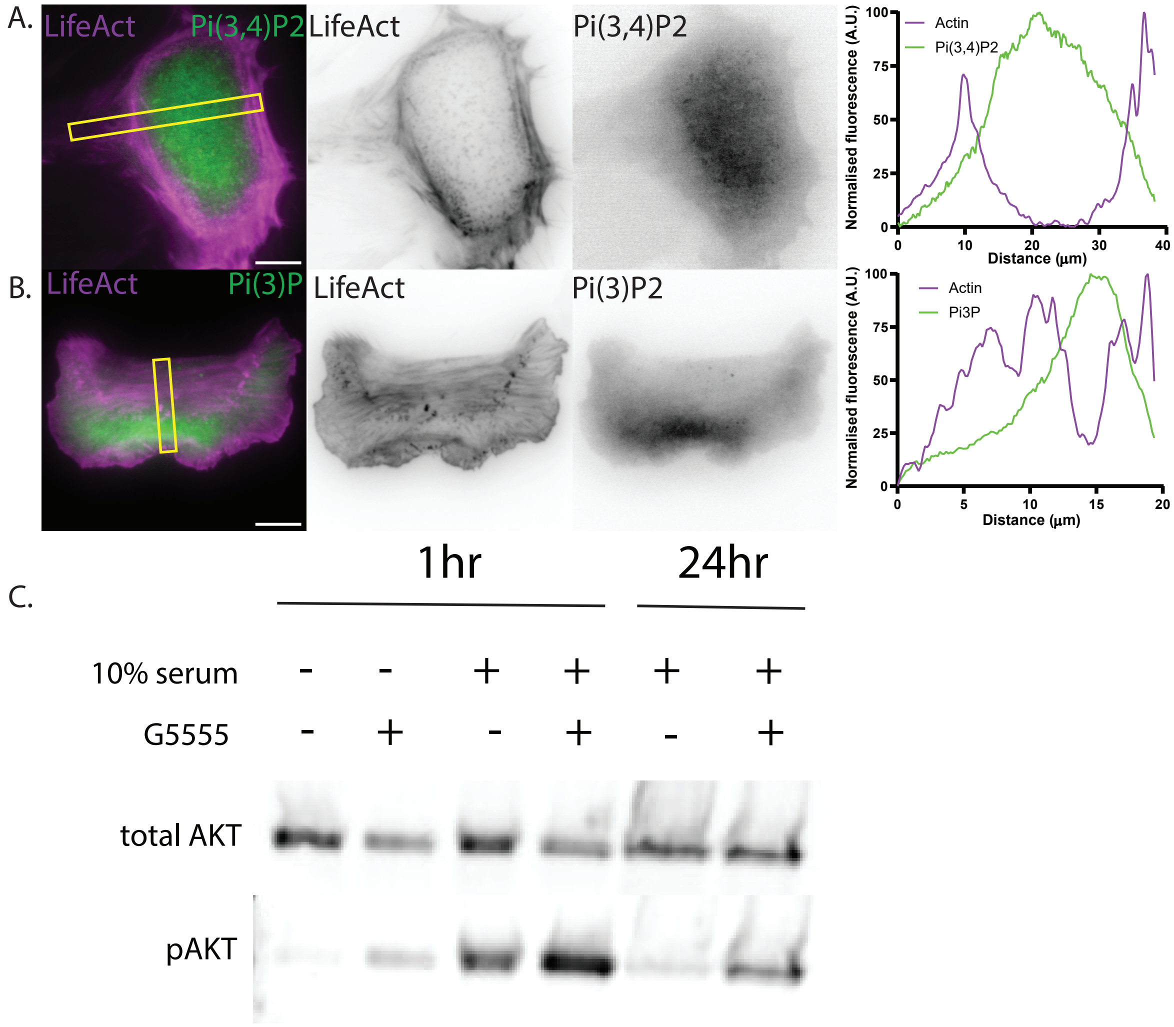


Supplimental Figure 4: Localisation of 3' phosphoinositide's and pAKT signalling. (A-B). Representative TIRFm image of cell expressing LifeAct (shown in magenta) and indicated fluorescently tagged phosphoinositide marker (shown in green), treated with G5555. Accompanying linescan highlights localisation of phosphoinositide marker to actin ring. (A.) MEF expressing GFP-PH-Tapp1 and mCherry-LifeAct. (B.) MEF expressing p40PX-EYFP and mCherry-LifeAct. (C.) Western blot showing change in pAKT levels in response to G5555 addition. All scale bars 10µm.

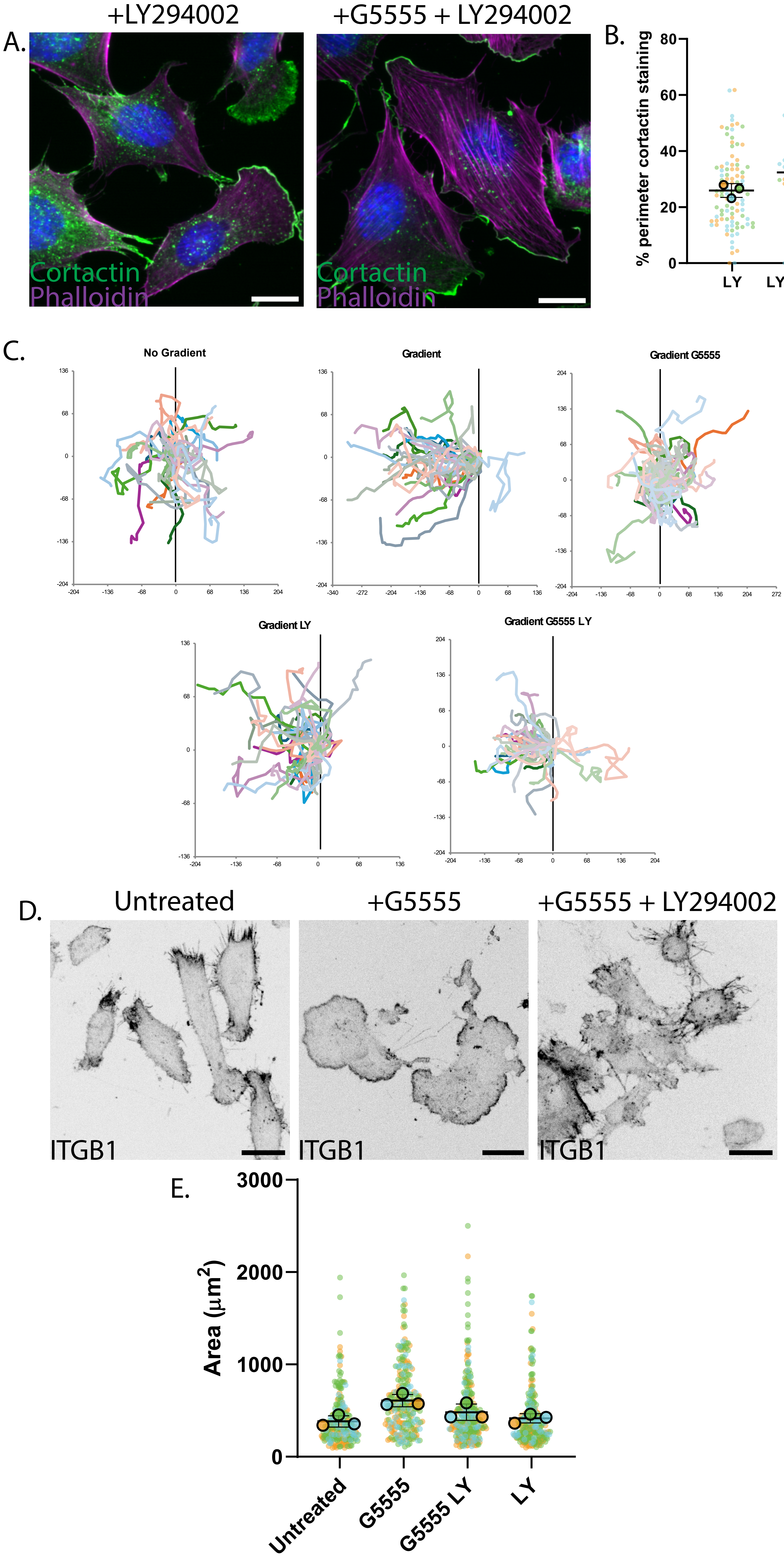


Supplemental Figure 5: PI3K dependence of phenotypes observed upon PAK kinase inhibition. (A.) Representative images showing cortactin localization in cells treated with LY294002 or cotreated with LY294002 and G5555 (B.) Quantification of the percentage of perimeter of cell marked by cortactin in cells treated with LY294002 or cotreated with LY294002 and G5555. (C.) Subset of tracks from cells migrating in indicated condition. Gradient indicates presence of a chemotactic gradient. (D.) Representative images of MiaPacca cells in indicated conditions. Cells were fixed and stained for ITGB1 to allow easy visualization. (E.) Quantification of MiaPacca cell area in indicated conditions. All scale bars 10μm.
